# Apical length governs computational diversity of L5 pyramidal neurons

**DOI:** 10.1101/754499

**Authors:** Alessandro R. Galloni, Aeron Laffere, Ede Rancz

## Abstract

Anatomical similarity across the neocortex has led to the common assumption that the circuitry is modular and performs stereotyped computations. Layer 5 pyramidal neurons (L5PNs) in particular are thought to be central to cortical computation because of their extensive arborisation and nonlinear dendritic operations. Here, we demonstrate that computations associated with dendritic Ca^2+^ plateaus in L5PNs vary substantially between the primary and secondary visual cortices. L5PNs in the secondary visual cortex show reduced dendritic excitability and smaller propensity for burst firing. This reduced excitability is correlated with shorter apical dendrites. Using numerical modelling, we uncover a universal principle underlying the influence of apical length on dendritic backpropagation and excitability, based on a Na^+^ channel-dependent broadening of backpropagating action potentials. In summary, we provide new insights into the modulation of dendritic excitability by apical dendrite length and show that the operational repertoire of L5 neurons is not universal throughout the brain.

## Introduction

The neocortex is thought to have a modular structure composed of ‘canonical circuits’ performing stereotyped computations (Harris & Shepherd, 2015; Markram et al., 2015; Miller, 2016). Anatomical evidence supports the existence of repeating circuit architectures that display similar general features across species and brain areas (Carlo & Stevens, 2013; Douglas & Martin, 2004; Mountcastle, 1997). It is generally thought that these architectural motifs serve as a physical substrate to perform a small range of specific, canonical computations (Bastos et al., 2012; Braganza & Beck, 2018).

Pyramidal neurons are the main building blocks of these circuit motifs. Across brain areas and species, their biophysical attributes endow them with non-linear properties that allow them to implement a repertoire of advanced computations at the single cell level (Gidon et al., 2020; London & Häusser, 2005; Spruston, 2008). Layer 5 pyramidal neurons (L5 PNs) in particular provide a striking example of how dendritic properties can underlie circuit-level computations in a laminar circuit. Their dendritic supralinearities enable signal amplification and coincidence detection of inputs — a crucial operation to integrate feedforward and feedback streams that often send projections onto separate dendritic domains. In these cells, a single backpropagating action potential (bAP), when combined with distal synaptic input, can trigger a burst of somatic action potentials. The crucial mechanism underlying this supralinear phenomenon is the all-or-none dendritic Ca^2+^ plateau (M. E. Larkum, Kaiser, & Sakmann, 1999; Matthew E. Larkum, Zhu, & Sakmann, 1999).

Morphology and intrinsic properties have a profound influence on neuronal excitability. Dendritic topology and the electrical coupling between the soma and dendrites is thought to be particularly crucial for determining a neuron’s integrative properties (Mainen & Sejnowski, 1996; Schaefer, Larkum, Sakmann, & Roth, 2003; van Ooyen, Duijnhouwer, Remme, & van Pelt, 2002; Vetter, Roth, & Hausser, 2001). Recent experimental work has shown that there can be substantial variation in intrinsic properties of L5 neurons depending on the location within a cortical area or on the species they are recorded from (Beaulieu-Laroche et al., 2018; Fletcher & Williams, 2019). However, it is often assumed that pyramidal neurons have robust enough properties across cortical areas and brain structures to support similar computations (Bastos et al., 2012; Hawkins, Ahmad, & Cui, 2017; M. Larkum, 2013; Shipp, 2016). For instance, analogous to L5 PNs, hippocampal pyramidal neurons also display dendritic Ca^2+^ APs that support coincidence detection of distal and proximal inputs (Jarsky, Roxin, Kath, & Spruston, 2005).

If L5 pyramidal neurons indeed have a common repertoire of operations in support of canonical computations, one would expect the same cell type in adjacent and closely related areas to exhibit the same computational repertoire. Here we have studied the bursting properties of thick-tufted L5 neurons (ttL5) in mouse primary and secondary visual cortices (V1 and V2m). Through systematic and rigorously standardized experiments, we found fundamentally different operation patterns linked to morphology in the two brain areas. Through computational modelling, we reveal new insights into biophysical mechanism linking excitability to morphology, which is able to account for this difference. Our results question the notion of a common operational repertoire in pyramidal neurons and thus cortical canonical computations as well.

## Results

### Thick-tufted L5 neurons in V2m lack BAC firing

We made whole-cell patch clamp recordings from ttL5 pyramidal neurons in V1 and V2m in acutely prepared mouse brain slices. To ensure consistency in cell type, recordings were restricted either to neurons projecting to the lateral posterior nucleus of thalamus, identified using retrograde labelling with cholera toxin subunit B (Supplementary Figure 1), or to neurons labelled in the Glt25d2-Cre mouse line (Groh et al., 2010). In addition, we were able to confirm the characteristic morphological features of ttL5 neurons in a subset of the recorded neurons using biocytin reconstructions. We were thus able to maintain cortical area as the primary variant when comparing V1 and V2m neurons.

To reproduce the conditions required for triggering BAC firing, we stimulated synaptic inputs near the distal tuft in L1 using an extracellular electrode in conjunction with somatic stimulation through the recording electrode (Fig. 1a). To avoid recruiting inhibitory inputs during the extracellular stimulation and create the most favourable conditions to enable BAC firing (Perez-Garci, Gassmann, Bettler, & Larkum, 2006), we added the competitive GABA_B_ receptor antagonist CGP52432 (1 µM) to the extracellular solution. Extracellular current pulses in L1 were adjusted to evoke either a subthreshold EPSP or a single action potential at the soma. Somatic injection of a 5 ms depolarizing current pulse through the recording electrode was used to trigger single APs. In V1 neurons, combined stimulation (with the L1 input triggered at the end of the somatic pulse) could evoke a prolonged plateau potential resulting in a burst of 3 APs. We repeated these experiments in ttL5 pyramidal neurons located in V2m under the same recording conditions. Upon coincident somatic AP and extracellular L1 stimulation, BAC firing was almost never observed in V2m, suggesting a much-reduced dendritic excitability in V2m neurons. For the purposes of these experiments, we defined as “supralinear” any cell in which three or more APs could be evoked following combined somatic and L1 stimulation (each evoking no more than one AP individually). Supralinearity was observed in half the recorded V1 neurons (10/20), while neurons in V2m showed an almost total lack of supralinearity (1/19, Fig. 1b).

**Figure 1.**
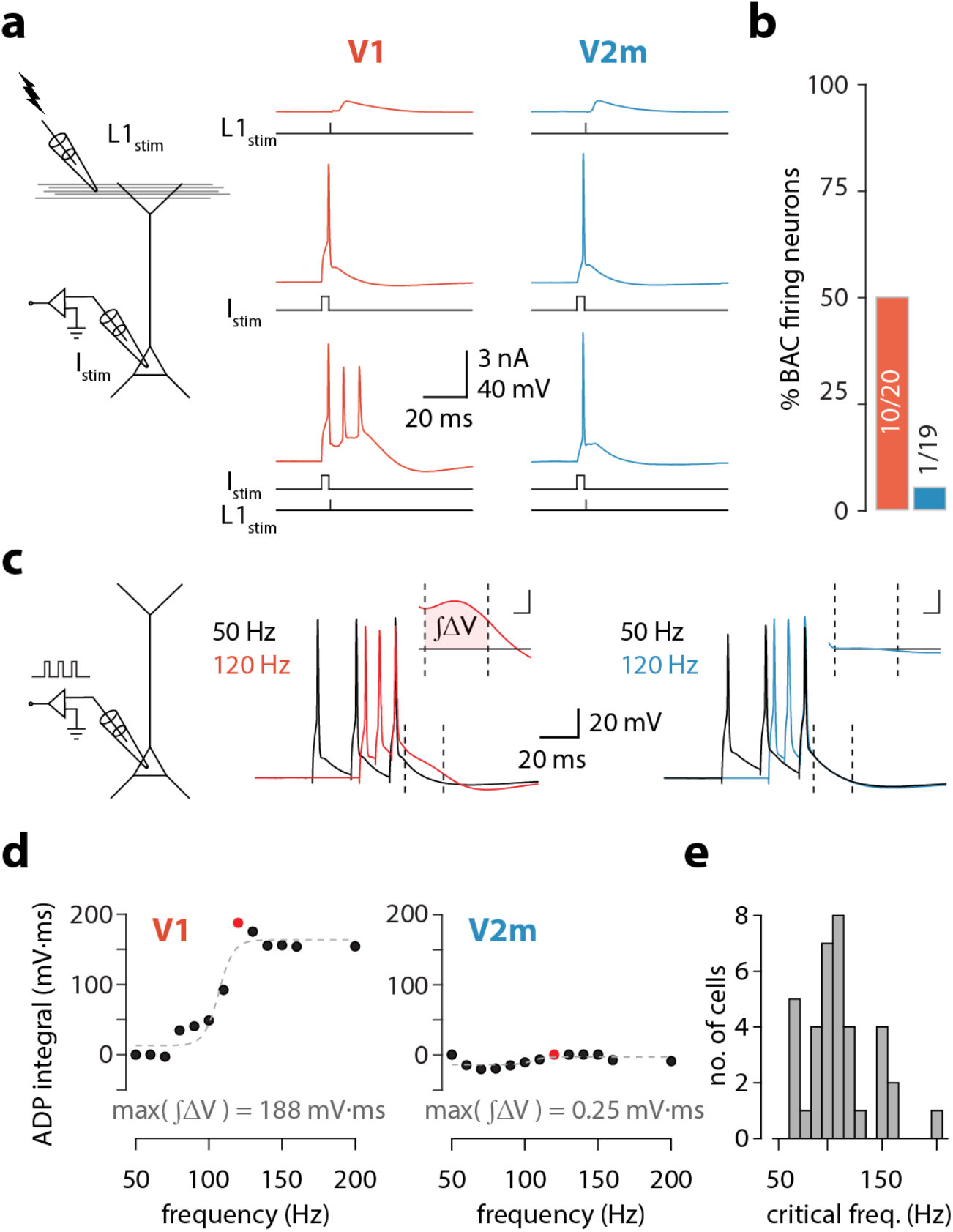
V2m neurons are less prone to burst than in V1. **a.** *Left:* diagram of experimental configuration. *Right:* example traces during BAC firing protocol, recorded from V1 (red) and V2m (blue) ttL5 neurons. **b.** Proportion of supralinear cells in V1 and V2m **c.** *Left:* diagram of experimental configuration. *Right:* example traces of V1 and V2m ttL5 neurons stimulated with 50 Hz and 120 Hz AP trains. Note the sustained after-depolarization following the 120 Hz spike train in the V1 neuron. *Inset:* ADP measured as the area between the 50 Hz trace and the higher frequency trace following the last spike. Inset scale bar: 5 ms x 5 mV. **d.** Quantification of ADP area at each measured frequency for the example neurons in *c*. The peak integral value is highlighted in red. **e.** Histogram of results across all recorded cells with defined critical frequency.

### Thick-tufted L5 neuron in V2m lack a critical frequency ADP

To further investigate the prevalence of dendritic supralinearities in ttL5 neurons across visual cortices, we recorded another hallmark of dendritic Ca^2+^ plateaus: a prominent somatic ADP following a high-frequency train of somatic APs (M. E. Larkum et al., 1999; Shai, Anastassiou, Larkum, & Koch, 2015). We recorded the somatic membrane potential from ttL5 neurons and evoked three action potentials using 3 ms pulses of somatic current injection at frequencies ranging from 50 Hz to 200 Hz in 10 Hz increments (Fig. 1c). In V1 neurons, increasing the AP frequency above a critical frequency typically resulted in a sudden increase in the ADP (Fig. 1c, **middle**). However, when recording in V2m under the same experimental conditions, there was usually no change in ADP, even at firing frequencies as high as 200 Hz (Fig. 1c, **right**). To quantify this effect, we aligned the peaks of the last AP for each frequency and measured the area of the ADP difference between the 50 Hz trace and the higher frequency traces in a 20 ms window (4–24 ms) following the last AP (Fig. 1c, **inset**). This measure of ADP increased sharply above a critical frequency and was often largest around the value of this frequency (Fig. 1d). The mean critical frequency across all cells in both V1 and V2m was 110.8 ± 29.6 Hz (SD, n = 37, excluding cells that did not have a critical frequency, Fig. 1e).

Next, we measured the maximal ADP integral value for each cell (Fig. 2a), regardless of the presence of a critical frequency. Neurons in V2m had significantly smaller ADP area (V1 mean = 91 ± 50 mV*ms, SD, n = 41; V2m mean = 42 ± 33 mV*ms, SD, n = 49; p = 4.54 * 10^−6^, two-sample Kolmogorov-Smirnov test), reflecting that most of these cells lacked a critical frequency altogether. The extracellular artificial cerebrospinal fluid (ACSF) contained either 1.5 or 2 mM CaCl_2_. As there was no statistically significant difference between the two conditions in either V1 or V2m (p > 0.05, two-sample Kolmogorov-Smirnov test; Supplementary Figure 2), we pooled all recordings. Similarly, as there was no significant difference in the ADP measure across V2m neurons labelled retrogradely or by the Glt25d2-Cre line (p = 0.617, two-sample Kolmogorov-Smirnov test, Supplementary Figure 3), we pooled these two populations. To obtain an unbiased count of cells showing supralinearity, we separated the unlabelled maximum ADP values pooled from both V1 and V2m into two groups using k-means clustering (with k = 2). N.B. that in this experiment the definition of supralinear classification differed from the experiments of Fig. 1a. The percentage of neurons classified as supralinear (summarized in Fig. 2b) was more than three times higher in V1 than in V2m, regardless of the specific supralinearity measure. In both the BAC firing and ADP experiments mentioned above, bursting was typically also apparent in the spiking response to a long (500 ms) depolarization at the soma. While all ttL5 neurons are generally characterized by a spike doublet at the beginning of the current step, in bursting neurons there is also a critical current step above which the initial spike burst is substantially larger, usually with 3 or 4 spikes followed by a deeper afterhyperpolarization (Supplementary Figure 4).

**Figure 2.**
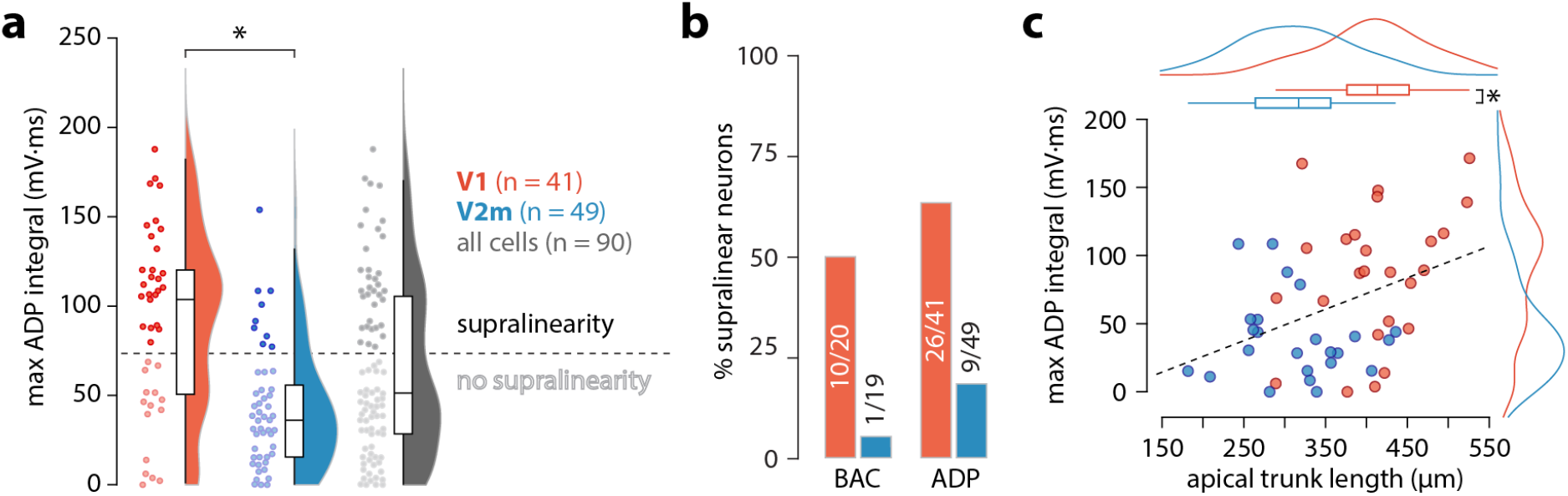
Excitability correlates with apical trunk length. **a.** Summary data of peak ADP integral values for all recorded neurons. The dashed line indicates the division between the two groups of cells classified through k-means clustering, drawn halfway between the cell with the lowest maximum ADP in the “supralinearity” cluster and the cell with the highest value in the “no supralinearity” cluster. **b.** Proportion of supralinear cells in V1 and V2m. **c.** Length of the apical trunk (soma to main bifurcation) plotted against the corresponding maximum ADP integral values. Dashed line is a linear fit; curves at the top and right are kernel density plots of the two variables in V1 and V2m.

These results show a much-diminished dendritic excitability, and as such different integrative properties, in V2m ttL5 neurons compared to V1 under the same conditions and in the same operational ranges. Previous research has indicated the length of the apical trunk as a possible factor involved in determining the dendritic excitability of ttL5 neurons in V1 (Fletcher & Williams, 2019). To test this, we reconstructed the apical trunk of 22 V1 and 26 V2m neurons from those recorded. Apical trunk lengths were significantly shorter in V2m than in V1 (V1 mean = 409 ± 64 µm, SD, n = 22; V2m mean = 313 ± 65 µm, SD, n = 26; p = 4.26*10^−6^, two-sample t-test, Fig. 2c). Additionally, there was a correlation between maximal ADP integral values and apical trunk length across the two populations (p = 5.37*10^−3^; F-test). These results suggest that there may be a surprisingly counter-intuitive interaction between apical trunk length and dendritic excitability—the longer the trunk, the more excitable the neuron.

### BAC firing is absent in short ttL5 models

To investigate possible mechanisms underlying the dependence of bursting on apical trunk length, we ran numerical simulations in conductance-based compartmental models of ttL5 neurons. We first probed BAC firing in a morphologically detailed model developed by Hay et al. (Hay, Hill, Schürmann, Markram, & Segev, 2011), using the model parameters (biophysical model 3) and morphology (cell #1) favoured for reproducing BAC firing. As in the original paper, BAC firing was triggered by injecting a 0.5 nA current at the apical bifurcation coupled to a somatic action potential evoked by square-pulse current injection at the soma. Mirroring the responses seen in the subset of strongly bursting ttL5 neurons, coincident stimulation triggered BAC firing in the detailed model (Fig. 3a, **left**). We then applied the same model to an example V2m morphology with a shorter apical dendrite. The Ca^2+^ channel hotspot was moved to the new apical branch point (350–450 µm from the soma vs 685-885 µm in the long morphology). The amplitude of the dendritic current injection (0.194 nA) was scaled so as to obtain the same depolarization amplitude at the bifurcation in both model cells. With this morphology, coincident tuft and somatic stimulation evoked only a single somatic spike and did not trigger a dendritic Ca^2+^ plateau (Fig. 3a, **right**). To ensure comparability between the long and short morphology, BAC firing was probed with both 100 µm and 200 µm Ca^2+^ channel hotspot size. We did not observe any qualitative effect of hotspot size in either model (Supplementary Figure 5). To test if Ca^2+^ plateaus were at all possible in the short neuron model, we stimulated the short neuron with a large current injection (0.5 nA) at the dendritic electrode. While the resulting dendritic potential was substantially larger, showing activation of calcium conductances, it resulted in only a small depolarization at the soma. Even when combining the large current injection with a somatic spike, no spike burst could be triggered (Supplementary Figure 6).

**Figure 3.**
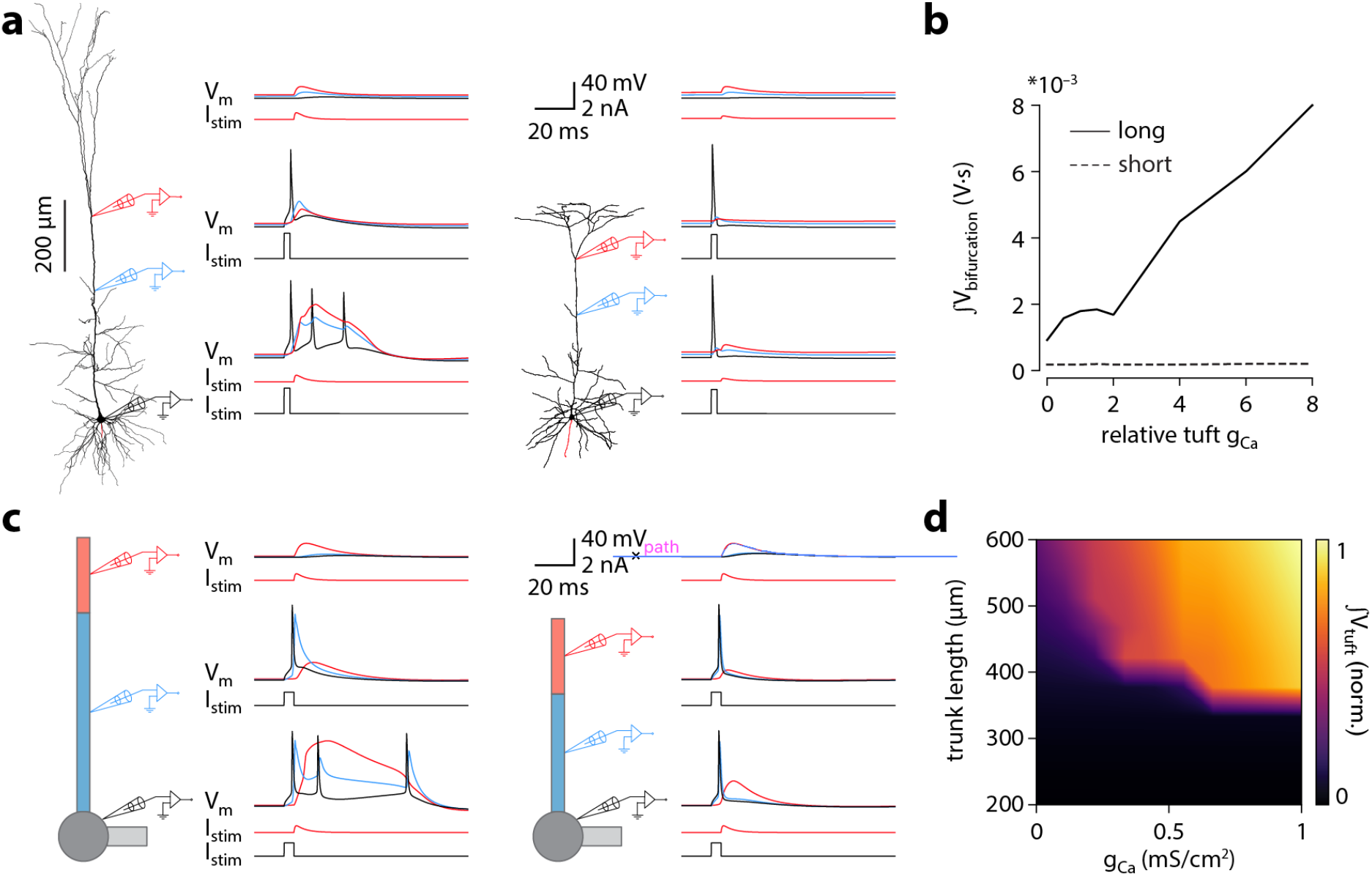
Shorter model neurons are less prone to burst. **a.** *Left:* detailed morphology of a ttL5 pyramidal neuron from the model favoured by Hay et al. for reproducing BAC firing. *Right:* reconstructed morphology of a ttL5 neuron recorded in V2m. Injected current and recorded voltage traces are shown for the soma (black), the apical trunk (blue, 400 and 200 µm), and the main bifurcation (red, 620 and 370 µm) under three different stimulation paradigms. **b.** Integral of voltage at the branch point during coincident somatic and branch point stimulation, plotted against relative g_Ca_. **c.** *Left:* diagram of the reduced neuron model. Apical trunk length 800 μm. Injected current and recorded voltage traces as in *a*. *Right:* Same for a version of the reduced model modified to have an apical trunk length of 200 μm. **d.** Heat map representing the normalised tuft voltage integral during combined somatic and tuft stimulation in the reduced model, plotted against the absolute density of Ca^2+^ channels in the tuft compartment and the length of the apical trunk compartment. Default g_Ca_ ≅ 0.45 mS/cm^2^.

To explore the sensitivity of Ca^2+^ plateaus to dendritic Ca^2+^ channel density in the long and short neurons, we scaled the sum Ca^2+^ conductance (g_Ca_) between 0 and 8 times the original values. To minimize the number of variables, when scaling the relative g_Ca_ we kept the ratio of the two channels (low- and high-voltage activated) constant. In the long morphology the integral of the distal dendritic voltage, acting as an indicator of the large and sustained depolarization during a Ca^2+^ plateau, increased proportionally to g_Ca_. In the short morphology, however, the voltage integral stayed constant across all g_Ca_ values (Fig. 3b). This indicates that, although the size of a Ca^2+^ plateau depends on g_Ca_ in long neurons, in short neurons there is no Ca^2+^ channel activation and the magnitude of the voltage integral therefore does not depend on g_Ca_.

To be able to vary dendritic length across a continuous range of values, we turned to a reduced ttL5 model based on Bahl et al. (Bahl, Stemmler, Herz, & Roth, 2012). The simplicity of this model has the added benefit of reducing the number of variables, allowing us to explore general principles of dendritic voltage propagation with more clarity. As with the morphologically detailed model, the reduced model with the original published parameters displayed BAC firing triggered by coincident tuft and somatic stimulation (Fig. 3c, **left**). Shortening the apical trunk was sufficient to eliminate this response (Fig. 3c, **right**).

We explored the dependence of BAC firing on apical trunk length and g_Ca_ by measuring the time-integral of tuft voltage as an indicator of Ca^2+^ plateau potentials (Fig. 3d). The presence of a Ca^2+^ plateau depended strongly on apical trunk length and was only sensitive to g_Ca_ above a critical length of approximately 350 µm (≅ 0.35 λ). Below this length, no Ca^2+^ plateaus were triggered regardless of how high g_Ca_ was set to. These experiments show that a reduced model can also reproduce our results, allowing us to explore and dissect the underlying parameters in more detail.

### Active propagation enhances voltage in long dendrites

To obtain a mechanistic understanding of what causes the length dependence of bursting, we made recordings from the final segment of the apical trunk as well as the tuft using the reduced model of Fig. 3c. To recreate the experimental conditions of Fig. 1b, we triggered 3 spikes at 100 Hz through a somatic electrode. As with coincident bAP and tuft input, increasing the length of the apical trunk facilitated dendritic Ca^2+^ plateau initiation (Fig. 4a). Interestingly, the width and peak voltage in the tuft increased steadily with dendritic length (Fig. 4b,c), even in the absence of Ca^2+^ currents (g_Ca_ = 0). In the presence of voltage-gated Ca^2+^ channels, the increased amplitude of bAPs triggered a large all-or-none Ca^2+^ plateau above a certain threshold length.

**Figure 4.**
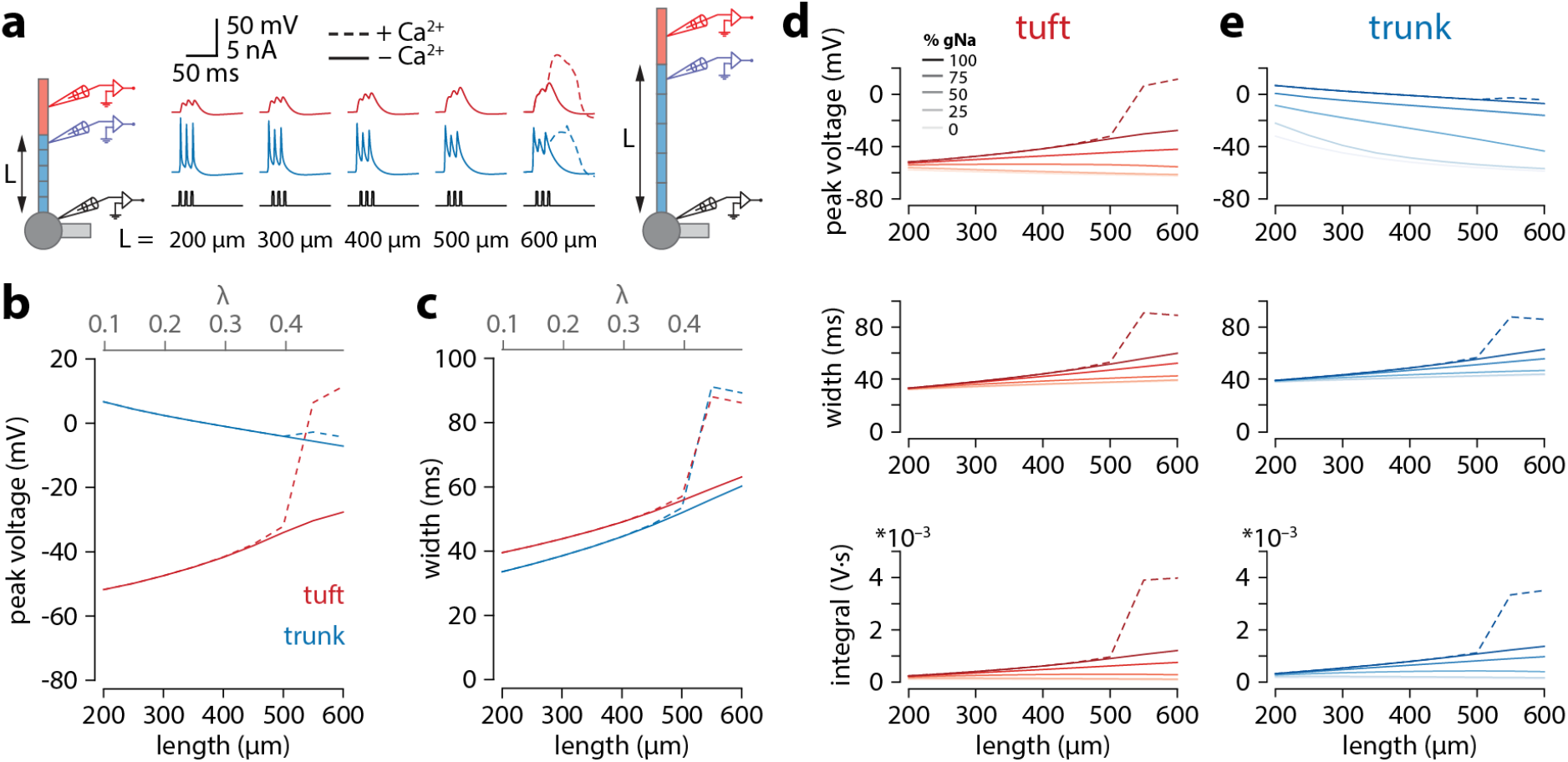
Tuft voltage increases with trunk length due to the widening of bAPs. **a.** Schematic of the simulation: stimulation site in the somatic compartment, and recording sites at the distal end of the apical trunk (blue) and in the centre of the tuft (red). Stimulus shown in black. Solid lines: g_Ca_ = 0; dashed lines: original g_Ca_. **b.** Peak voltage reached in trunk (blue) and tuft (red) for a range of simulations with different trunk lengths. Length constant λ = 1009 µm. **c.** Same as in *b* but plotting the width of the depolarization measured 2 mV above baseline. **d,e.** Peak voltage, width and integral values measured in the trunk and tuft for dendrites containing different Na^+^ channel densities in the apical trunk.

We found that bAP amplitude in the tuft increased as a function of apical trunk length despite a decreasing bAP amplitude in the distal segment of the trunk (Fig. 4b). Conversely, the width of the bAP (measured 2 mV above baseline) increased in both the tuft and trunk with length (Fig. 4c). While waveform broadening is a natural consequence of passive filtering along dendrites, the sustained voltage in the distal trunk required active dendritic propagation. In the reduced model, this active propagation in the apical trunk was mediated primarily by voltage-gated Na^+^ channels. Removing these channels caused a substantial reduction in peak voltage and width of the depolarization in the distal trunk, and importantly also abolished the trend of increasing tuft voltages with longer dendritic trunks (Fig. 4d,e). More generally, active propagation caused bAPs to be larger and broader at all distances along a long dendrite compared to the same absolute distances in shorter dendrites (Supplementary Figure 7). Consequently, when comparing the final positions along the trunk, the peak voltage is only marginally smaller in long dendrites despite the larger distance from the soma. This is not the case in a passive dendrite, where voltage attenuation depends on distance alone and is not sensitive to trunk length. We next tested how the specific distribution of active conductances affected the results. When all conductances were uniformly distributed along the apical trunk, the waveforms did not substantially change, and the enhanced voltage continued to trigger Ca^+^ plateaus only in neurons with long apical trunks (Supplementary Figure 8). We have also tested the specific contribution of the H-current (I_h_). Reducing I_h_ in the tuft had no effect on the length dependence of excitability. Reducing I_h_ in both the trunk and tuft resulted in the same qualitative finding, having only a marginal quantitative effect (Supplementary Figure 9). The general phenomenon of enhanced voltage propagation in longer dendrites resulting in amplification of tuft voltage thus did not depend on any of the particular model parameters above.

While it might seem counterintuitive that peak tuft voltage is increasing when the trunk voltage is decreasing, we propose that the temporal broadening of the depolarization can at least partially account for this via a passive mechanism. Wider depolarizations allow the tuft compartment to charge to a higher voltage. The rate and peak value of tuft charging depends on the passive properties of the tuft. The peak value of depolarization and the rate of voltage change are proportional to membrane resistance (R_m_) and membrane capacitance (C_m_), respectively. The product of these two parameters gives the membrane time constant (*τ*). To illustrate this, we applied voltage-clamp to the end of the distal segment of the trunk and delivered 30 mV square voltage pulses of increasing width (Fig. 5a). Due to capacitive filtering, short voltage steps did not fully charge the tuft while wide voltage steps allowed voltage to reach the steady-state values commanded by R_m_. To directly test the hypothesis that the relationship between trunk depolarization width and tuft membrane time constant caused the bAP amplitude in the tuft to increase with length, one could vary R_m_ by changing g_leak_. However, this would affect resting membrane potential and consequently alter voltage-dependent properties in the tuft. In order not to affect other variables in the model, we therefore chose to vary C_m_. For a given value of depolarization amplitude and width, increasing C_m_ (and therefore *τ*) in the tuft caused a reduction in the peak tuft voltage (Fig. 5b). These simulations show that the tuft time constant and the width of the bAP interact to create a higher tuft depolarization with longer apical trunks.

**Figure 5.**
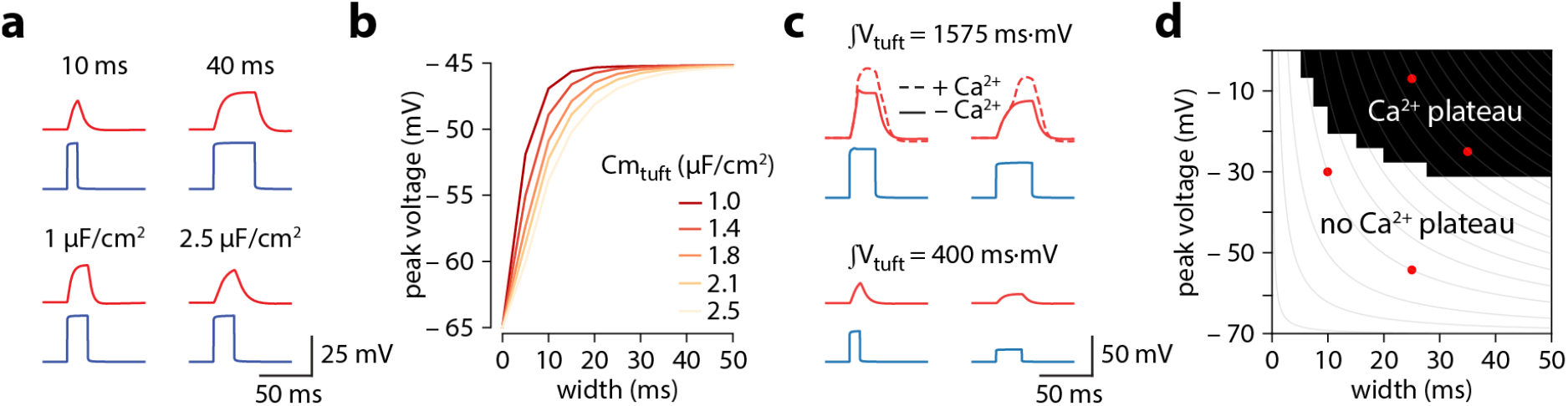
Tuft voltage increases with trunk length due to the widening of bAPs. **a.** Different width voltage steps injected into the trunk (blue) under differing tuft membrane capacitance conditions. Recorded tuft voltage in red. **b.** Peak voltage values reached in the tuft for a range of trunk step widths and tuft capacitances. In the original model, tuft C_m_ ≅ 1.75 µF/cm^2^. **c.** Same as in *a* but showing voltage steps of different width and amplitude with the same voltage integral in the trunk. For large integrals (top), the tuft voltage crosses the threshold for a Ca^2+^ plateau, while for smaller integrals (bottom) the voltage remains subthreshold. **d.** Voltage and width combinations for square voltage steps in the distal trunk which result in a Ca^2+^ plateau in the tuft. Red points represent the values used in *h.* Grey lines indicate width and amplitude combinations with equal integral.

It has previously been suggested that axial resistance (R_a_) in the apical dendrite may influence the backpropagation efficiency in dendrites and burstiness of ttL5 neurons (Fletcher & Williams, 2019). To test this hypothesis, we measured peak voltage and width in the trunk and tuft for different trunk lengths under different R_a_ conditions. We found that peak tuft voltage (and therefore burstiness) increased with increasing trunk R_a_, reaching the highest voltage near the reduced model’s original value, and decreasing again for higher values (Supplementary Figure 10). However, in these simulations burstiness always increased with trunk length regardless of R_a_. This indicates that, although important, R_a_ was not the primary determinant for generating the length-dependent effect and if R_a_ indeed correlates with length, these effects may combine to further enhance the tuft voltage in long neurons.

Overall, the combination of increased width and a relatively small reduction in amplitude resulted in a trunk voltage integral that increased with trunk length, thereby passing more charge to the adjacent tuft compartment. However, if active backpropagation was reduced or absent, the trunk integral and resulting tuft voltage decreased with length (**Fig.4d,e**). The peak tuft voltage approximately followed the integral of voltage in the distal trunk. To illustrate this, we applied voltage-clamp to the end of the trunk and injected square steps with a range of integrals obtained through various combinations of width and amplitude (Fig. 5c). This revealed a zone above a critical trunk integral for which many different width and depolarization combinations were sufficient to evoke a Ca^2+^ plateau in the tuft (Fig. 5d).

## Discussion

We made whole-cell patch-clamp recordings from layer 5 pyramidal neurons in primary and secondary visual cortices of mice. We found that both BAC firing and critical frequency ADP were almost entirely absent in the V2m neurons. Moreover, we observed that excitability was positively correlated with the length of the apical dendrite trunk across all neurons. To investigate the influence of apical trunk length on burstiness, we ran numerical simulations in compartmental biophysical models. Both morphologically detailed and reduced models showed that decreasing the apical trunk length resulted in reduced dendritic excitability. Further simulations revealed that this is due to an interplay between bAP width, amplitude and tuft impedance that depends critically on the presence of voltage-gated Na^+^ channels in the apical trunk. These results show that the same cell type, in closely related and adjacent cortical areas, and under the same operating conditions can have a very different computational repertoire. Our findings are thus inconsistent with the notion of canonical computations at the single cell level, suggesting they may not exist at the circuit level either.

Contrary to common assumptions, we observed considerable differences in the properties of ttL5 neurons across different brain regions. BAC firing, a dendritic operation, and critical frequency ADP, a measure of dendritic excitability, which are both critically dependent on dendritic Ca^2+^ plateaus, were almost completely absent in V2m. One key factor known to control dendritic excitability is inhibition (Perez-Garci et al., 2006). To determine the role of inhibition on the observed differences in excitability, we pharmacologically blocked GABA_B_ receptors, thus creating more favourable conditions for BAC firing. To make comparisons rigorous, we have used the same experimental protocols and conditions across all experiments. Furthermore, to exclude bias in selecting ttL5 neurons, we have recorded only from two well-defined groups, identified either by their projection target or genetic label. We found similarly diminished excitability in both groups of V2m neurons. The extracellular stimulation used to evoke BAC firing could in principle recruit different polysynaptic circuits in V1 and V2m, which could account for the difference. However, this would not explain the differences measured with the critical frequency paradigm using intracellular stimulation only. Another confounding variable affecting excitability could be the Ca^2+^ concentration in the extracellular solution. We have found the same results under different physiologically relevant concentrations. Altogether, these results show that decreased excitability of V2m ttL5 neurons is a robust phenomenon.

Correlated with the differences in excitability, we found ttL5 neurons in V2m to have significantly shorter apical trunks compared to V1 neurons. This data is consistent with recent structural MRI data (Fletcher & Williams, 2019) showing a thinner cortical mantle in more medial and posterior parts of the cortex. Using an existing widely used biophysical model designed to reproduce ttL5 properties such as BAC firing (Hay et al., 2011), we found that the same model applied to a shorter morphology resulted in a loss of BAC firing, independently of Ca^2+^ channel density. This was also true in a reduced ttL5 model with a simplified morphological structure (Bahl et al., 2012), which allowed for continuous exploration of apical trunk length. We found a sharp cut-off at a length of 0.35 λ (≅ 350 µm in model space), below which no BAC firing could be evoked. We note, however, that the reduced model is based on rat neurons and the apical trunk and oblique dendrites are pooled into the same compartment. It is also worth noting that, as the reduced model does not have a distinct compartment to represent the apical bifurcation, all Ca^2+^ channels are placed in the tuft compartment. Therefore, the numerical values of model length do not translate directly into apical trunk lengths for real mouse neurons.

Through our simulations, we identified voltage-gated Na^+^ channels in the apical dendrite as a key factor for reproducing our results. The Na^+^ channels control dendritic excitability by supporting active backpropagation, resulting in reduced attenuation over distance. Combined with a broadening of the bAP proportionally to trunk length (due to capacitive filtering), this caused longer neurons to have larger voltage integrals in the distal trunk, leading to greater charging of the tuft. Indeed, the peak tuft voltage depended on the amount of passive charging it experienced, which was determined by the membrane time constant in the tuft as well as by bAP width and amplitude. We hypothesised that, above a minimal threshold for peak trunk voltage, the primary determinant of peak tuft voltage is the time-averaged voltage in the trunk. Supporting this view, we found that the overall voltage integral was more important for triggering Ca^2+^ plateaus than the particular combination of depolarization width and amplitude.

Aside from dendritic length, our results do not exclude the involvement of other mechanisms to modulate excitability. For example, differences in axial resistance could influence the way voltage propagates along dendrites. While axial resistance indeed had a marked effect on the backpropagation of action potentials in our models, this was independent of trunk length and thus cannot account for our finding. Variations in density and activation properties of other voltage-gated channels may also influence dendritic excitability. It is interesting to note that, in the presence of Na^+^ channels, the bAP in neurons with longer trunks was larger and broader at every distance from the soma. This may be due to a cooperative effect of each trunk section on the sections both up- and downstream, with the voltage at each location decaying slower because of the more depolarised state of the remaining dendrite. Our data thus predicts that bAP width and amplitude measured at the same absolute distance from the soma differ across neurons with different apical trunk lengths.

There are a few notable counterexamples to the principle observed here. For example, the human ttL5 neuron was recently shown to have greater compartmentalization and reduced excitability compared to rat neurons despite being substantially longer (Beaulieu-Laroche et al., 2018). This may still be consistent with our predictions, as human ttL5 neurons also had reduced ion channel densities, which we show to be crucial for the length-dependent enhancement. Furthermore, as the boosting effect of a broader depolarization is subject to saturation (when the depolarization is wide enough to fully charge the tuft), we would expect the positive effect of length on tuft voltage to not increase monotonically. Consequently, beyond a certain length the trunk voltage would attenuate to the point where it is no longer sufficient to trigger a Ca^2+^ plateau. On the other end of the spectrum, layer 2/3 pyramidal neurons in rat barrel cortex have shorter apical trunks yet do show critical frequency ADP, although they do not exhibit spike bursts during BAC firing (M. E. Larkum, Waters, Sakmann, & Helmchen, 2007). It remains to be seen if the principle of length-dependent excitability generalizes to other species, or across more widespread cortical areas and cell types.

There may be important clinical implications to gaining a better understanding of how variations in cortical thickness, and the resulting changes in neuronal morphology, affect the physiology and computational properties of pyramidal neurons. Indeed, altered cortical thickness has been implicated in several debilitating neurological diseases and mental health conditions. For example, cortical thinning is used as a biomarker for Alzheimer’s disease (Dickerson et al., 2009) and correlates strongly with bipolar disorder (Hanford, Nazarov, Hall, & Sassi, 2016), while increased cortical thickness is present during development in individuals with autism (Khundrakpam, Lewis, Kostopoulos, Carbonell, & Evans, 2017). Interestingly, altered dendritic excitability has also been strongly implicated in several of these diseased conditions (Hall et al., 2015; Nanou & Catterall, 2018; Spratt et al., 2019). We uncovered a possible mechanistic link between cortical thickness and excitability, highlighting a new potential avenue of study for understanding the pathophysiology in these conditions and raising the prospect of identifying intervention targets.

Our findings on dendritic excitability in ttL5 neurons have wide-ranging implications for cortical computation. Feedback connectivity between cortical areas tends to target superficial layers while feedforward input favours basal dendrites (Coogan & Burkhalter, 1990; Rockland & Pandya, 1979). BAC firing is believed to play a major role in integrating these two pathways to modulate sensory perception (Takahashi, Oertner, Hegemann, & Larkum, 2016) and to enable brain-wide learning algorithms that would otherwise be intractable (Guerguiev, Lillicrap, & Richards, 2017; Sacramento, Ponte Costa, Bengio, & Senn, 2018). This is particularly relevant in higher order cortical areas, which are more likely to process brain-wide feedback to integrate convergent multisensory, motor and cognitive signals (Freedman & Ibos, 2018). However, we show that, at least within the secondary visual cortex, different operations must be implementing the multimodal integration of top-down and bottom-up signals.

Our findings thus challenge the commonly held notion that the neocortex is composed of canonical circuits performing stereotyped computations on different sets of inputs across the brain (Douglas & Martin, 2004; Harris & Shepherd, 2015; Markram et al., 2015; Mountcastle, 1997). The heterogeneity of operating modes may expand the ability of cortical areas to specialize in the computations that are required for processing their particular set of inputs, at the cost of reduced flexibility in generalizing to other types of input. It may also imply that the non-linear operations performed through Ca^2+^ plateaus and BAC firing may not be required outside primary sensory cortices, perhaps because in these regions the input hierarchy is less defined. It may therefore be more important to maintain equal weighting between different sensory modalities and rely on other mechanisms to change the weights according to the reliability of each input. With such simple morphological adjustments capable of generating a wide range of possible operations, the brain undoubtedly leverages the array of available computations to improve cognitive processing.

## Materials and Methods

### Animals

All animal experiments were prospectively approved by the local ethics panel of the Francis Crick Institute (previously National Institute for Medical Research) and the UK Home Office under the Animals (Scientific Procedures) Act 1986 (PPL: 70/8935). Wild-type and transgenic male mice were used. Tg (Colgalt2-Cre)NF107Gsat (MGI:5311719, referred to as Glt) and Tg (Rbp4-Cre)KL100Gsat (MGI:4367067, referred to as Rbp4) lines created through the Gensat project (Gerfen, Paletzki, & Heintz, 2013; Groh et al., 2010) were crossed with the Ai14 reporter line expressing tdTomato (MGI:3809524). Animals were housed in individually ventilated cages under a normal 12-hour light/dark cycle.

### Surgical procedures

Surgeries were performed on mice aged 3–7 weeks using aseptic technique under isoflurane (2–4%) anaesthesia. Following induction of anaesthesia, animals were subcutaneously injected with a mixture of meloxicam (2 mg/kg) and buprenorphine (0.1 mg/kg). During surgery, the animals were head-fixed in a stereotactic frame and a small hole (0.5–0.7 mm) was drilled in the bone above the injection site. Alexa Fluor 488-conjugated Cholera toxin subunit B (CTB, 0.8% w/v, Invitrogen) was injected using a glass pipette with a Nanoject II (Drummond Scientific) delivery system at a rate of 0.4 nL/s. Injections of 100– 200 nL were targeted to the lateral posterior (LP) thalamic nucleus, with stereotactic coordinates: 2.2–2.5 mm posterior to bregma, 1.45 lateral of the sagittal suture, 2.45 mm deep from the cortical surface. To reduce backflow, the pipette was left in the brain approximately 5 minutes after completion of the injection before being slowly retracted.

### Slice preparation

Male mice (6–12 weeks old) were deeply anaesthetised with isoflurane and decapitated. In mice that were injected with CTB, this occurred at least 3 weeks after the injection. The brain was rapidly removed and placed in oxygenated ice-cold slicing ACSF containing (in mM): 125 sucrose, 62.5 NaCl, 2.5 KCl, 1.25 NaH_2_PO_4_, 26 NaHCO_3_, 2 MgCl_2_, 1 CaCl_2_, 25 dextrose; osmolarity 340–350 mOsm. Coronal slices (300 µm thick) containing visual cortex were prepared using a vibrating blade microtome (Leica VT1200S or Campden 7000smz-2). Slices were immediately transferred to a submerged holding chamber with regular ACSF containing (in mM): 125 NaCl, 2.5 KCl, 1.25 NaH_2_PO_4_, 26 NaHCO_3_, 1 MgCl_2_, 1.5 or 2 CaCl_2_, 25 dextrose; osmolarity 308–312 mOsm. The holding chamber was held in a water bath at 35 °C for the first 30–60 min after slicing and was kept at room temperature (22 °C) for the remaining time (up to 12 hours) after that. All solutions and chambers were continuously bubbled with carbogen (95% O_2_ / 5% CO_2_).

### Electrophysiology

After the 35 °C incubation period, individual slices were transferred from the holding chamber to the recording chamber, where they were perfused at a rate of ~6 mL/min with regular ACSF (see above) continuously bubbled with carbogen and heated to 35 ± 1 ̊C. Borosilicate thick-walled glass recording electrodes (3-6 MΩ) were filled with intracellular solution containing (in mM): 115 CH_3_KO_3_S, 5 NaCl, 3 MgCl_2_, 10 HEPES, 0.05 EGTA, 3 Na_2_ATP, 0.4 NaGTP, 5 K_2_-phosphocreatine, 0.5% w/v biocytin hydrochloride (Sigma), 50 µM Alexa Fluor 488 hydrazide (Invitrogen); osmolarity 290-295 mOsm; pH 7.3. Visually guided whole-cell patch-clamp recordings were targeted to neurons in L5 of medial V2 (V2m) and V1 that were fluorescently labelled with either CTB or with tdTomato (for Glt25d2-Cre mice), to ensure that the recordings were restricted to ttL5 neurons. Visual areas were defined based on approximate stereotactic coordinates (Franklin & Paxinos, 2007). All recordings were made in current-clamp mode. Extracellular monopolar stimulation was achieved by passing a DC current pulse (0.1-1 ms, 20-320 µA) through a glass patch-clamp pipette with a broken tip (~20 µm diameter) using a constant current stimulator (Digitimer DS3). Current was passed between two silver / silver chloride (Ag/AgCl) wires: one inside the pipette, which was filled with recording ACSF, and the other coiled around the outside of the pipette. In experiments using extracellular stimulation, 1 µM CGP52432 was added to the ACSF.

### Immunohistochemistry and morphological reconstructions

After recording, slices were fixed overnight at 4 °C in a 4% formaldehyde solution and were subsequently kept in PBS. For immunohistochemical detection, the fixed slices were first incubated for 1-2 hours at room temperature in blocking solution containing 0.5% Triton X-100 and 5% Normal Goat Serum (NGS) in PBS. Slices were then washed twice (10 min each) in PBS and incubated overnight in a staining solution containing 0.05% Triton X-100, 0.5% NGS, DyLight 594-conjugated streptavidin (2 µg/ml). Slices were then washed in PBS (3 times, 5 min each) and stained with DAPI (5 µg/ml) for 10 min. After another wash (3 times, 5 min each), slices were mounted on glass slides and images were acquired with a confocal microscope (Leica SP5, objective: 20x/0.7NA or 10x/0.4NA, pinhole size: 1 airy unit). The images were used to reconstruct the neurons with Neurolucida 360 (MBF bioscience).

### Data acquisition and analysis

Recorded signals were amplified and low-pass filtered through an 8 kHz Bessel filter using a MultiClamp 700B amplifier (Molecular Devices). Filtered signals were then digitized at 20 kHz with a National Instruments DAQ board (PCIe-6323). Acquisition and stimulus protocols were generated in Igor Pro with the Neuromatic software package (Rothman & Silver, 2018). Further analysis and data visualization were performed with custom macros and scripts written in Igor Pro and Matlab (MathWorks). Raincloud plots (consisting of a scatter plot, a box plot, and a kernel density plot) were generated in MATLAB using published scripts (Allen, Poggiali, Whitaker, Marshall, & Kievit, 2019). All box plots presented show the median, interquartile range, 2nd and 98th percentile of the dataset. Confocal images were processed with Fiji (https://fiji.sc/).

### Modelling

Simulations were performed with the NEURON (Carnevale & Hines, 2006) simulation environment (7.7.1) embedded in Python 3.6. To model the consequences of morphological differences between V1 and V2m ttL5 cells, we utilised existing models of ttL5 pyramidal cells with either accurate morphological detail (biophysical model 3, cell #1 from Hay et al. 2011 (Hay et al., 2011), referred to as detailed model) or simplified multicompartment morphologies (Ca^2+^ enriched model 2 from Bahl et al. 2012 (Bahl et al., 2012), referred to as reduced model). To study the effect of morphology in the detailed model, biophysical model 3 from Hay et al. 2011 (Hay et al., 2011) was applied to the reconstructed morphology from one of our recorded ttL5 neurons in V2m (which has a substantially shorter apical trunk than the morphology used in the original model). Each morphology contained low-voltage-activated (LVA) and high-voltage-activated (HVA) Ca^2+^ channels located in a 100-200 µm long region around the main apical bifurcation.

Subsequent simulations using the reduced model were done by modifying only selected parameters described in the results, such as the length of the apical trunk compartment, leaving all other parameters unchanged. Briefly, this reduced model (Bahl et al., 2012) is divided into sections representing the soma, axon (hillock and initial segment, AIS), basal dendrites, apical trunk, and apical tuft. Active conductances are present in all compartments and include the following: hyperpolarization-activated cation (HCN) channels (basal dendrite, apical trunk, tuft), transient voltage-activated Na^+^ (Na_t_) channels (soma, axon hillock, AIS, apical trunk, tuft), persistent voltage-activated Na^+^ (Na_p_) channels (soma), fast voltage-activated K^+^ (K_fast_) channels (soma, apical trunk, tuft), slow voltage-activated K^+^ (K_slow_) channels (soma, apical trunk, tuft), muscarinic K^+^ (K_m_) channels (soma), slow Ca^2+^ (Ca_s_) channels (tuft), Ca^2+^ dependent K^+^ (K_Ca_) channels (tuft), and a Ca^2+^ pump (tuft). The density of the K_fast_ and K_slow_ channels decays exponentially from the soma to the tuft. The density of Na_t_ channels decays linearly from the soma to the tuft, while HCN channels linearly increase in density. N.B. the tuft, but not the trunk, contains Ca^2+^ channels; consequently, there is no hotspot similar to the apical bifurcation in the detailed model. When varying trunk length, Na_t_, K_fast_, K_slow_, and HCN conductances in each trunk segment were redistributed so as to take into account the new distance of each segment from the soma (thereby changing the total conductance in the trunk). All code will be available on GitHub after acceptance.

## Acknowledgements

We thank Arnd Roth and Alexandra Tran-Van-Minh for advice on modelling. We are grateful to Lee Fletcher, Arnd Roth, Alexandra Tran-Van-Minh, Michael Hausser and Anna Cappellini for helpful comments on the manuscript.

## Author contributions

A.R.G.: Designed and performed biological experiments, designed modelling experiments, analysed data, wrote the paper. A.L.: Designed and performed modelling experiments, analysed data. E.A.R.: Conceptualized research, designed experiments, acquired funding, wrote the paper.

All authors discussed the results and implications and commented on the manuscript at all stages. There are no competing interests.

All authors declare no financial or non-financial competing interest.

## Supplementary figures

**Supplementary Figure 1.**
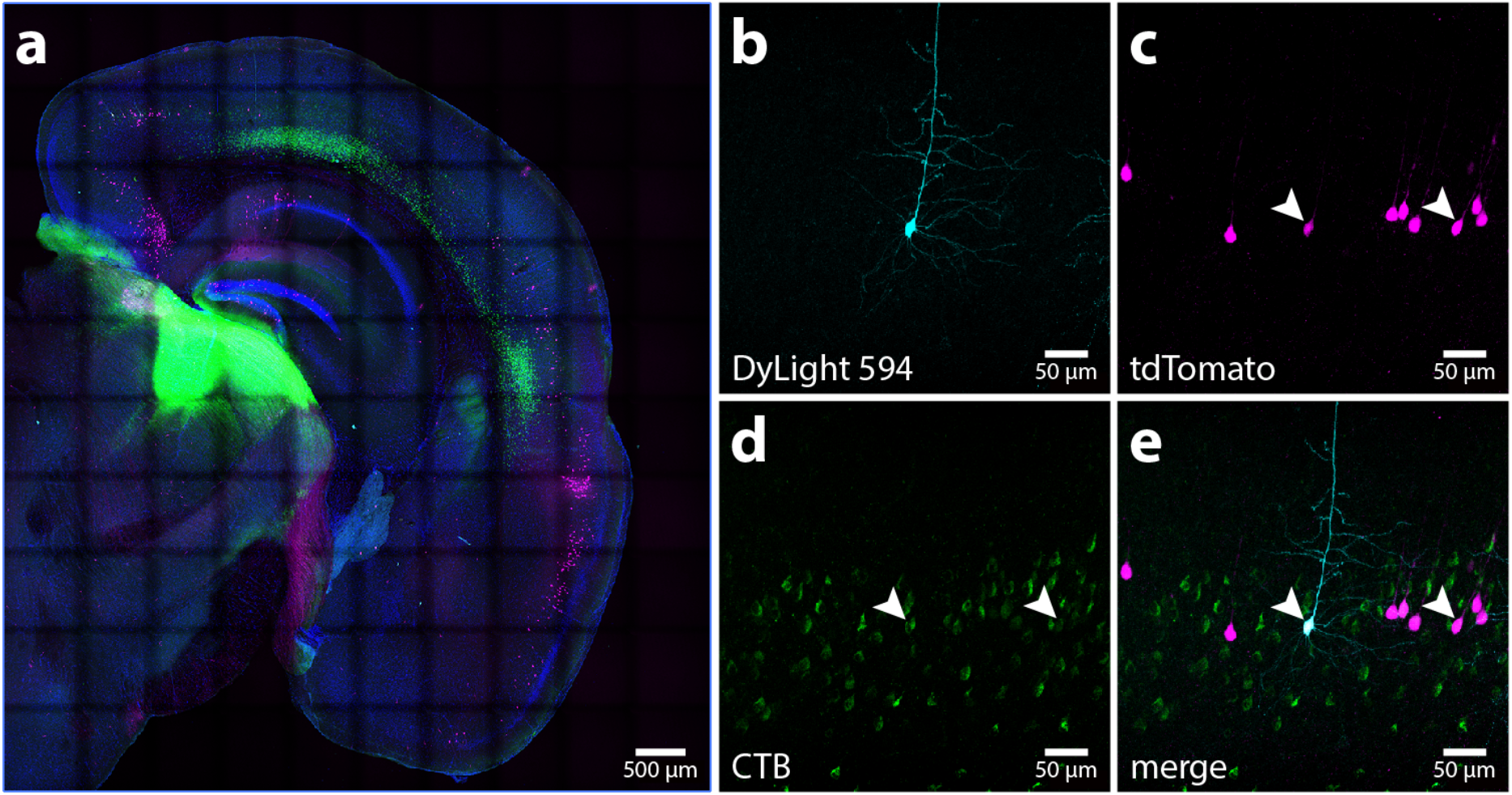
Confocal images from a Glt25d2-Cre mouse injected with CTB-Alexa Fluor 488 in LP. **a.** Coronal slice of one hemisphere containing visual cortex, showing the injection site and retrogradely labelled neurons (green), tdTomato-expressing Glt25d2-Cre neurons (magenta), a DAPI stain (blue), and neurons that have been filled with biocytin during intracellular recordings and stained with DyLight 594 (cyan). **b.** Biocytin-filled ttL5 neuron in V2m. **c.** Neighbouring tdTomato-expressing Glt25d2-Cre neurons. **d.** CTB-labelled L5 neurons projecting to LP. **e.** Composite image of the above. Arrowheads highlight two example cells labelled by both tdTomato and CTB.

**Supplementary Figure 2.**
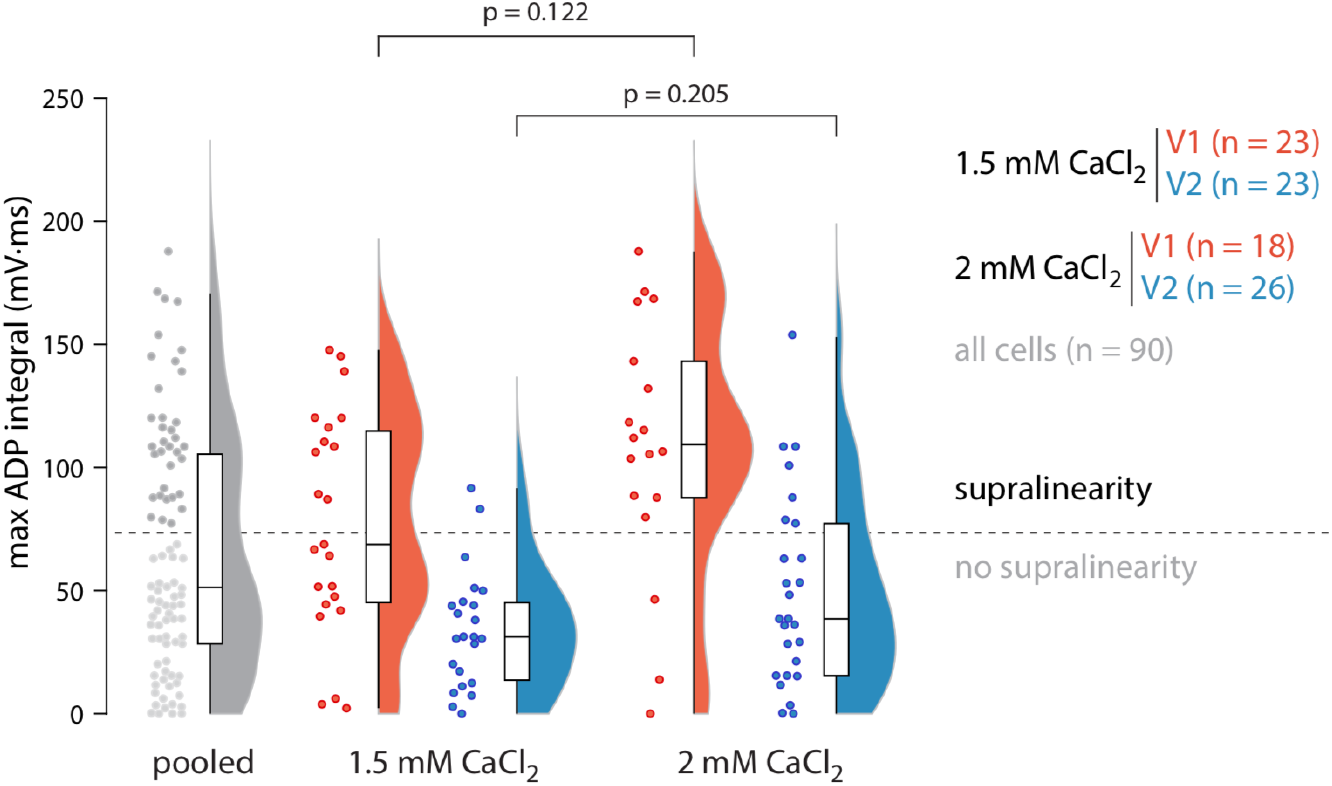
Maximum ADP integral for all cells split by recording ACSF containing either 1.5 or 2 mM CaCl_2_. p values for two-sample Kolmogorov-Smirnov test.

**Supplementary Figure 3.**
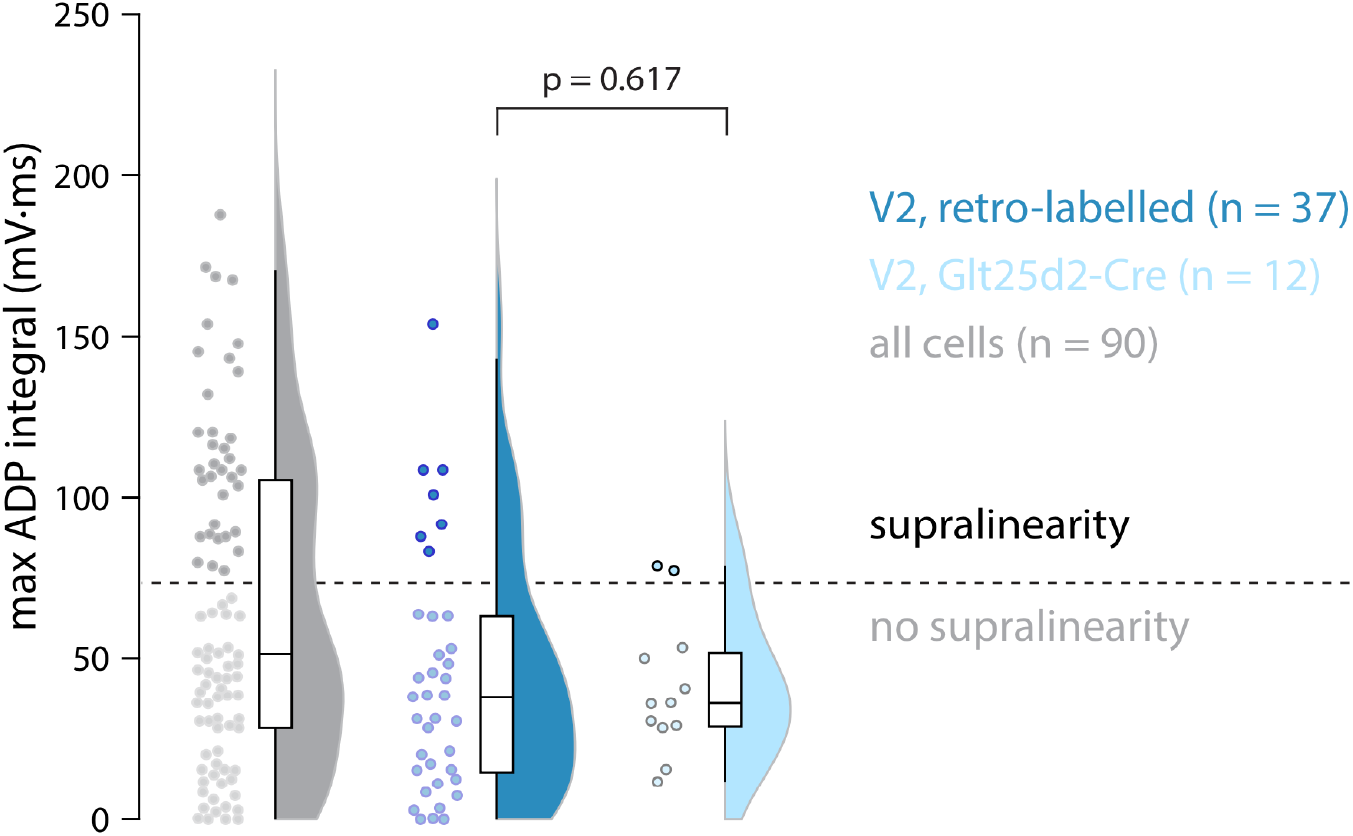
Maximum ADP integral for all cells including V1 (grey), retro-labelled cells recorded from V2m (dark blue), and V2m cells labelled in the Gltd2-Cre mouse line (light blue). p value for two-sample Kolmogorov-Smirnov test.

**Supplementary Figure 4.**
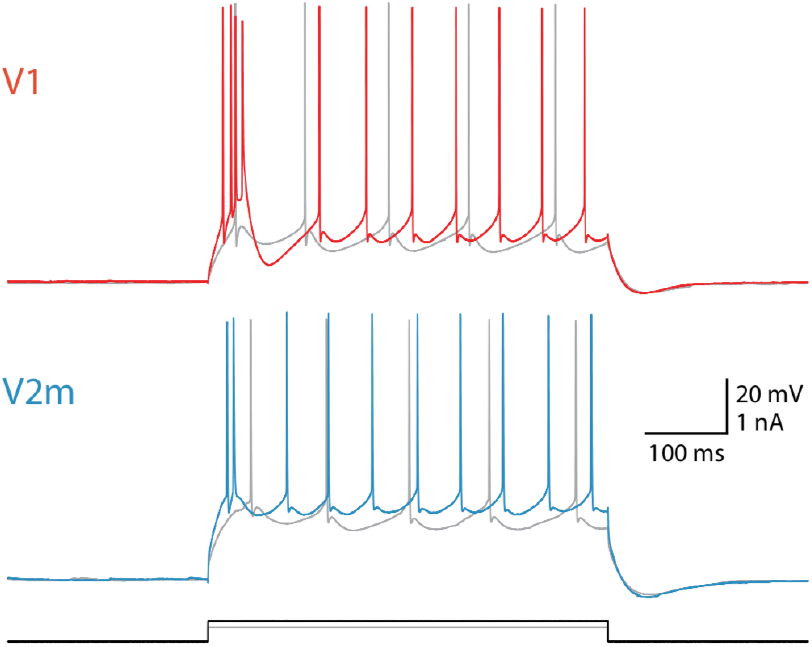
Representative voltage traces for V1 and V2m neurons in response to 500 ms wide depolarizing current steps. Stimulation at 60 pA (grey) and 180 pA (coloured or black) above rheobase (200 pA for both).

**Supplementary Figure 5.**
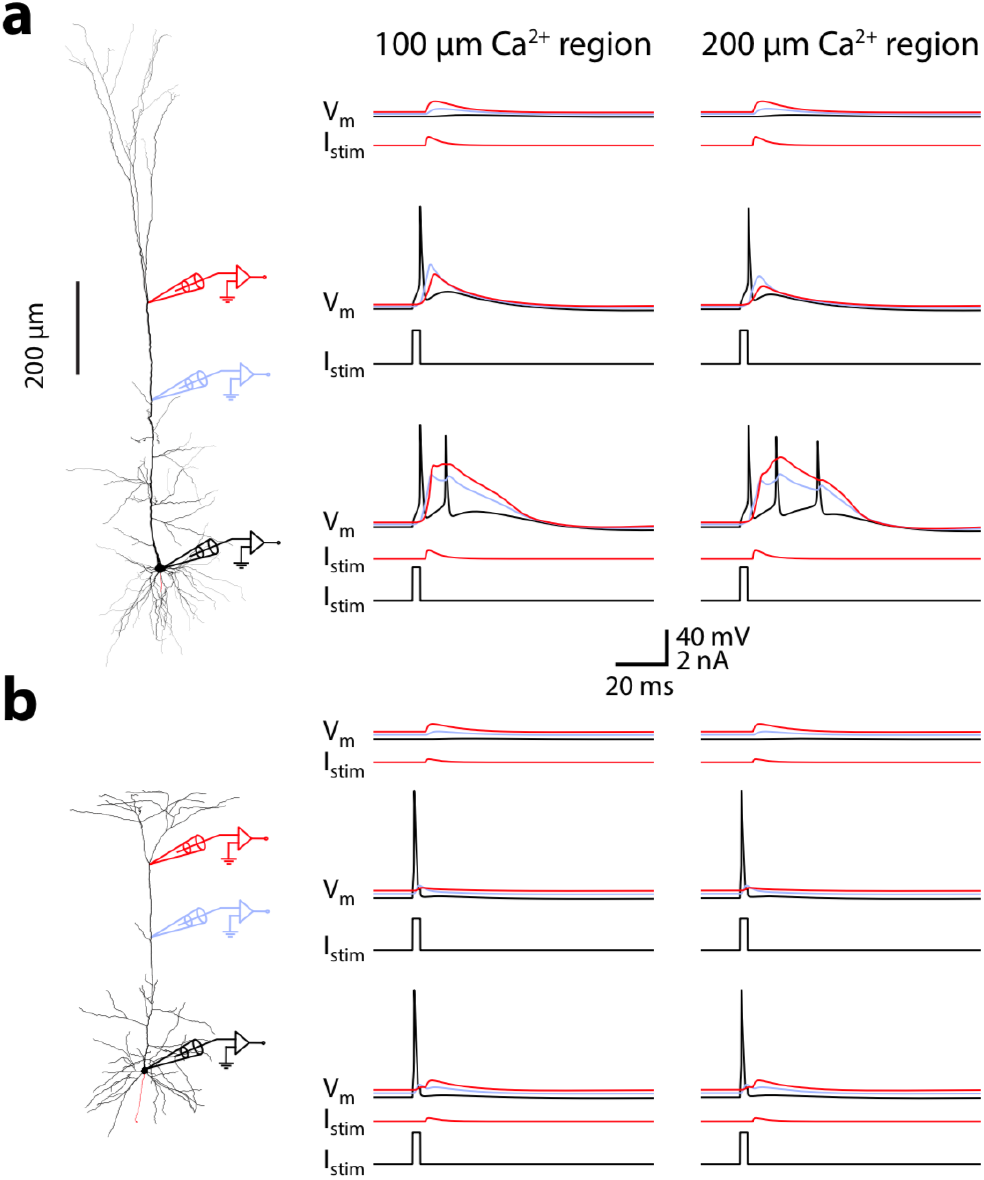
Ca^2+^ hot-spot size does not affect excitability. **a.** Voltage traces recorded from different parts of the long neuronal morphology containing either a 200 µm (as in the original), or 100 µm long Ca^2+^ region in the distal apical trunk. **b.** Same as in a, but for the shorter morphology.

**Supplementary Figure 6.**
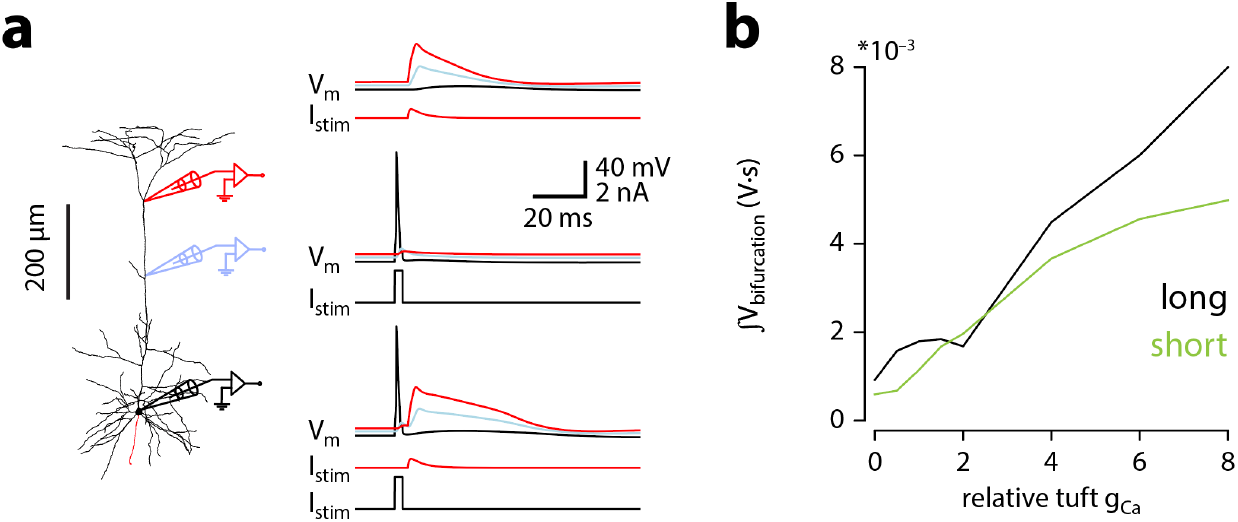
Large current injections to the nexus of short model cells can trigger calcium plateaus. Same as in *Fig. 3a & b*, but for the short morphology using the same dendritic current injection as in the long morphology (0.5 nA).

**Supplementary Figure 7.**
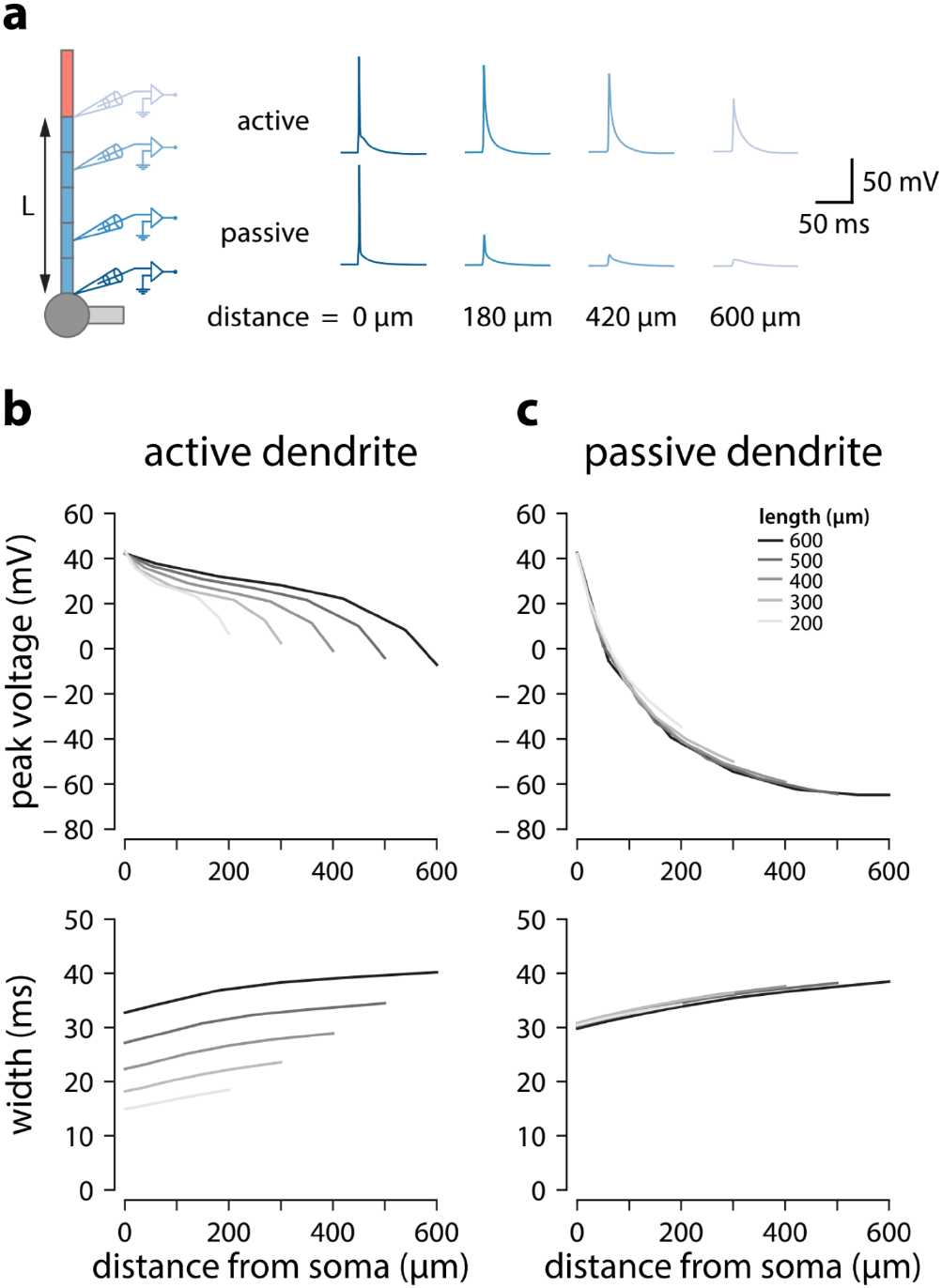
Backpropagation of APs in active and passive trunks of different length. **a.** Backpropagation of a somatic spike elicited through a single 3 ms wide 2 nA square current step at the soma in a model neuron with 600 µm apical trunk length. Voltage recordings at different distances along the trunk. **b.** Peak voltage and width measured at different absolute distances (same relative) for active model neurons. Width was measured as the interval between the voltage values 2 mV above baseline membrane potential. Colours indicate models with different apical trunk lengths. N.B. At any given absolute distance from the soma, peak voltage and width of the bAP are larger when the apical trunk is longer. **c.** Same as in *b* but with all voltage-dependent conductances removed from the trunk and tuft compartments. N.B. voltage attenuation is independent of trunk length.

**Supplementary Figure 8.**
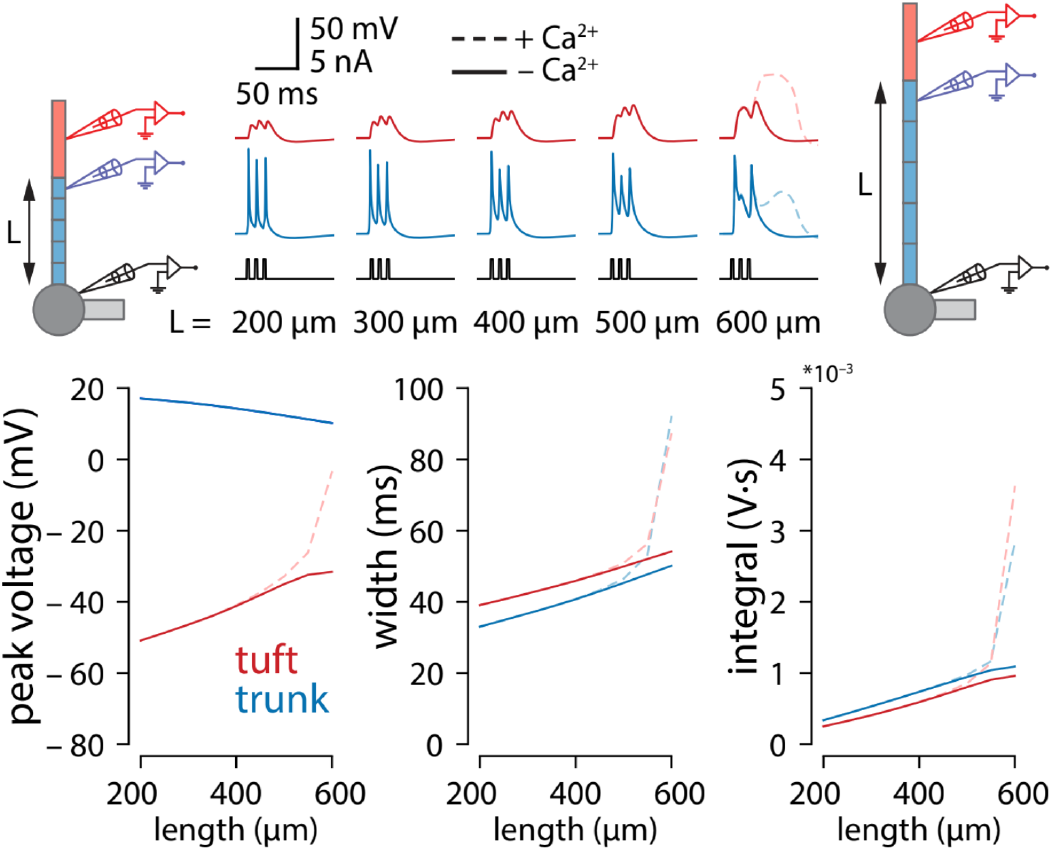
Tuft voltage increases with trunk length independently of conductance gradients. Same experiments as in Fig. 3a but with uniform distribution of all active conductances in the apical trunk. Total conductance was maintained for each channel.

**Supplementary Figure 9.**
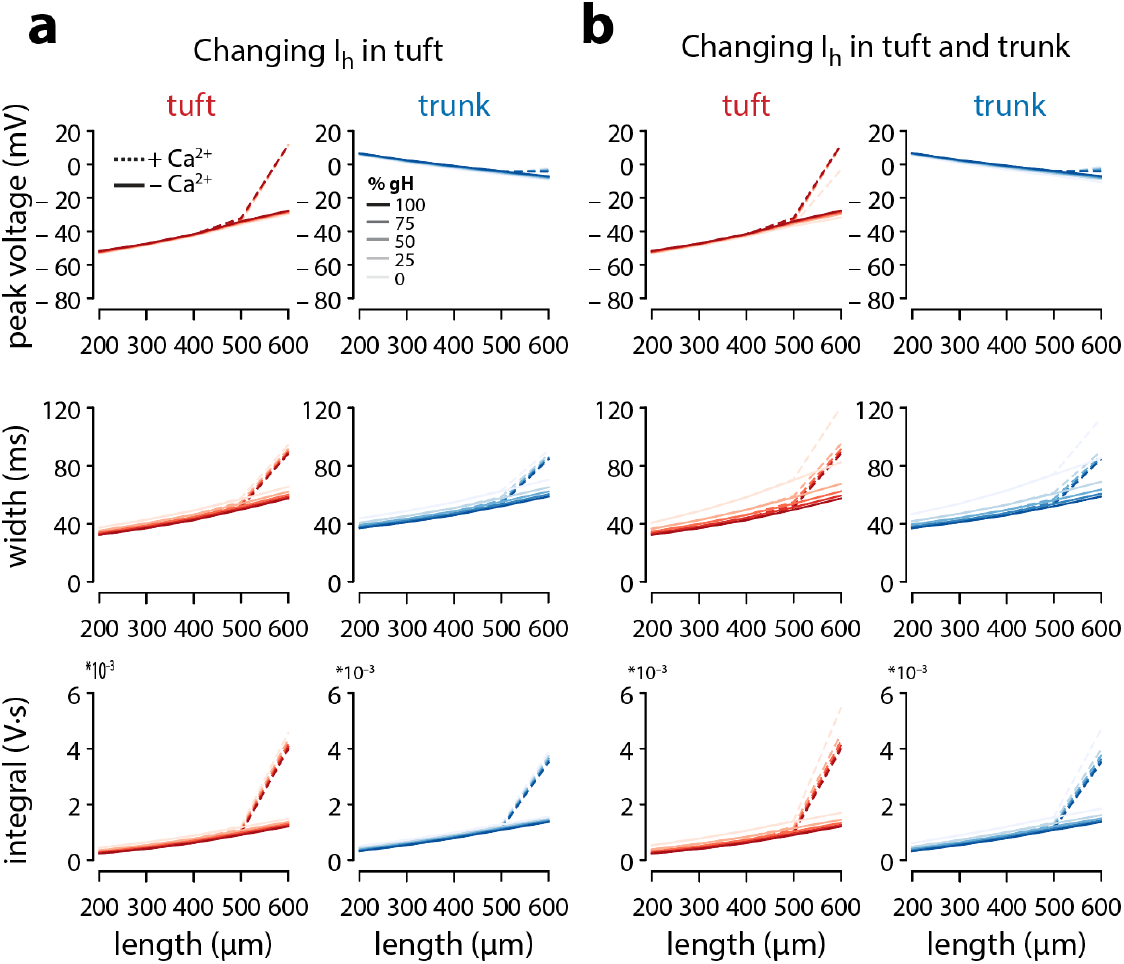
I_h_ does not contribute to length dependent increase of excitability. **a.** Plots of peak voltage, width and integral of bAPs under different g_HCN_ (shades) in the tuft, both in the presence (solid) or absence (dashed) of Ca^2+^ channels. **b.** Same as *a*, but varying g_HCN_ in both tuft and trunk compartments.

**Supplementary Figure 10.**
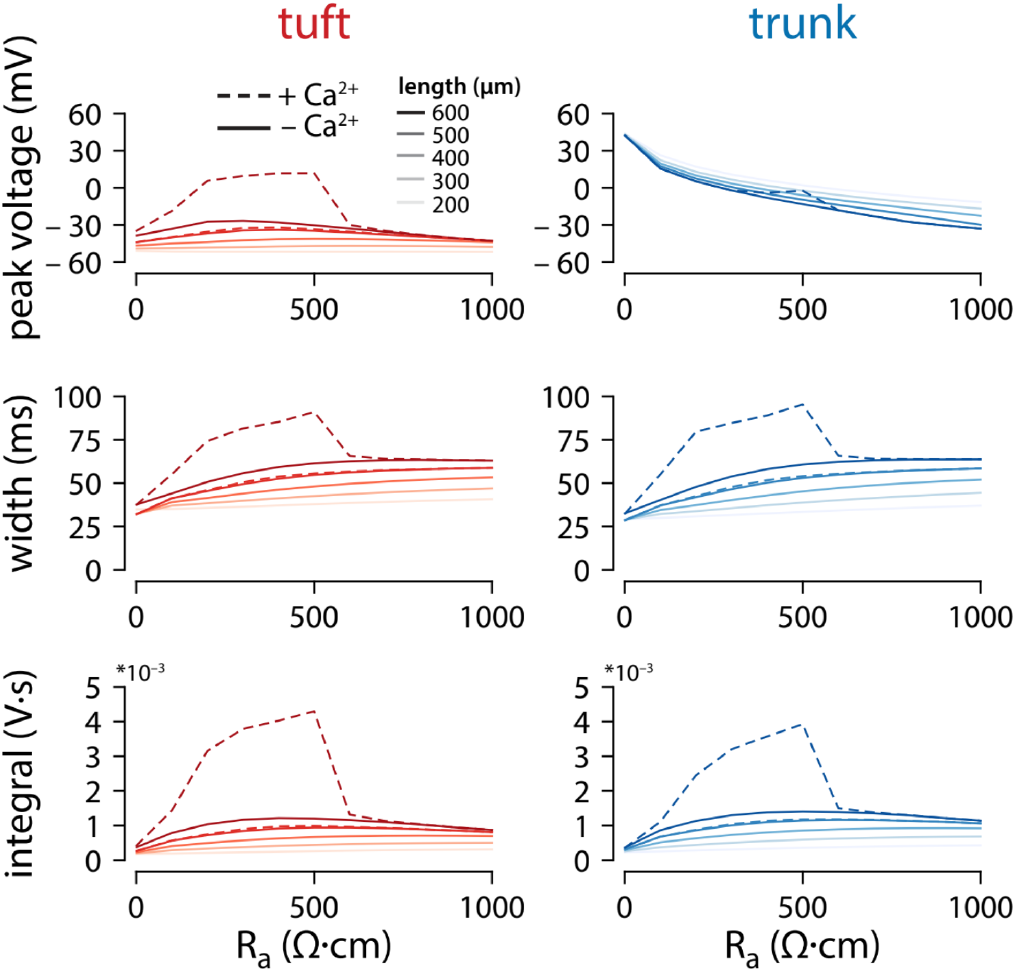
Effect of axial resistance on voltage propagation. Plots of peak voltage, width, and voltage integral reached in the tuft and trunk for varying values of trunk length and axial resistance (R_a_). Default R_a_ ≅ 382.22 Ω∙cm. The stimulus and recording conditions were the same as in Fig. 2a, with 3 APs at 100 Hz triggered in the somatic compartment. Solid lines show simulations with g_Ca_ = 0, while dashed lines show the same simulations with the original gCa = 0.45 mS/cm^2^.

